# Fine Mapping Regulatory Variants by Characterizing Native CpG Methylation with Nanopore Long-Read Sequencing

**DOI:** 10.1101/2024.09.27.614715

**Authors:** Yijun Tian, Shannon K. McDonnell, Lang Wu, Nicholas B. Larson, Liang Wang

**Affiliations:** Department of Tumor Microenvironment and Metastasis, Moffitt Cancer Center, Tampa, FL 33612, United States; Division of Clinical Trials and Biostatistics, Department of Quantitative Health Sciences, Mayo Clinic, Rochester, MN 55905, United States; Division of Cancer Epidemiology, Population Sciences in the Pacific Program, University of Hawai□i Cancer Center, University of Hawai□i at Mānoa, Honolulu, HI 96813, United States

**Author notes:** **Correspondence to:** Liang Wang, Department of Tumor Microenvironment and Metastasis, H. Lee Moffitt Cancer Center & Research Institute, Vincent Stabile Research Building 22403, 12902 Magnolia Drive, Tampa 33612, USA.

## Abstract

5-methylcytosine (5mC) is the most common chemical modification occurring on the CpG sites across the human genome. Bisulfite conversion combined with short-read whole genome sequencing can capture and quantify the modification at single nucleotide resolution. However, the PCR amplification process could lead to duplicative methylation patterns and introduce 5mC detection bias. Additionally, the limited read length also restricts co-methylation analysis between distant CpG sites. The bisulfite conversion process presents a significant challenge for detecting variant-specific methylation due to the destruction of allele information in the sequencing reads. To address these issues, we sought to characterize the human methylation profiling with the nanopore long-read sequencing, aiming to demonstrate its potential for long-range co-methylation analysis with native modification call and intact allele information retained. In this regard, we first analyzed the nanopore demo data in the adaptive sampling sequencing run targeting all human CpG islands. We applied the linkage disequilibrium (LD) R^2^ to calculate the co-methylation in nanopore data, and further identified 27,875, 50,481, 26,542 and 51,189 methylation haplotype blocks (MHB) in COLO829, COLO829BL, HCC1395 and HCC1395BL cell lines, respectively. Interestingly, while we found that majority of the co-methylation were in a short range (≤200bp), a small portion (1∼3%) showed long distance (≥1,000bp), suggesting potential remote regulatory mechanisms across the genome. To further characterize the epigenetic changes related to transcription factor binding, we profiled the 5mC percentage changes surrounding various motif sites in JASPAR collection and found that CTCF and KLF5 binding sites showed reduced methylation, while FOXE1 and ZNF354A sites showed increased methylation. To further investigate the allele-specific 5mCG in the prostate genome, we designed a target region covering methylation quantitative trait loci (mQTL) and genome-wide association study (GWAS) risk germline variants and generated long reads with adaptive sampling run in the 22Rv1 cell line. To identify the allele-specific methylation in the 22Rv1 cell line, we performed long-read based phasing and compared the 5mCG signals between the two haplotypes. As a result, we identified 6,390 haplotype-specific methylated regions in the 22Rv1 cell line (p-MWU ≤ 1e-5 and delta ≥ 50%). By examining haplotype-specific methylated regions near the phasing variants, we identified examples of allele-specific methylated regions that showed allele-specific accessibility in the ATAC-seq data. By further integrating the ATAC-seq data of 22Rv1, we found that methylation levels were negatively correlated with chromatin accessibility at the genome-wide scale. Our study has revealed native methylome profiling while preserving haplotype information, offering a novel approach to uncovering the regulatory mechanisms of the human prostate genome.

## Introduction

Over the past decades, enormous phenotypical connections have been identified between single nucleotide polymorphism (SNP) and complex disease risk in the Genome-Wide Association Study (GWAS). To help understand the underpinning biological meaning, the expression Quantitative Trait Locus (eQTL) analysis had been used to determine variants associated with the RNA expression of susceptible genes. Beyond the productivity of these association findings, challenges arise regarding prioritizing the causal (regulatory) variants and the target genes, which are not easily recognizable due to the linkage disequilibrium (Consortium, 2024; Nelson et al., 2024; Tian et al., 2023; Tian et al., 2022). The methylation Quantitative Trait Locus (mQTL) analysis is more mechanistically oriented, focusing on detecting associations between genetic variation and DNA methylation levels. This approach inherently suggests that DNA sequence variations may directly influence the DNA base modification of methylation sites, offering deeper insight into the regulatory role of SNPs. To distinguish between methylated and unmethylated cytosines, the bisulfite conversion is commonly used with microarray or next-generation sequencing to profile the feature at the genome-wide scale. However, due to the technical limitations, the conversion-based approach presents several challenges for methylation detection and mQTL interpretation. First and foremost, the conversion eliminates allele information for SNPs containing cytosine, which hampers haplotype identification for each sequencing read.

Additionally, when measuring co-methylation profiles, next-generation sequencing (NGS) methods face difficulties in evaluating distant CpG sites, primarily due to read length constraints. Moreover, PCR amplification during NGS library preparation can result in duplicated methylation patterns and introduce biases in 5mC detection. For microarray-based methylation profiling, only 1.5% to 3.0% of CpG sites in the human genome are covered, with the data provided as a summarized percentage rather than as precise base modification calls. Importantly, the existing mQTL experimental design depends on associations across samples between the genotypes of millions of SNPs and the methylation levels of thousands of CpG sites. This approach generates numerous correlations influenced by allele linkage disequilibrium (LD), leading to mQTL findings that are broadly distributed throughout the genome. Consequently, these findings may not be particularly useful for prioritizing functional variants or regulatory elements, especially in robust large-scale studies involving over 1,000 samples. In 4,170 whole-blood samples, Huan et al. (Huan et al., 2019) identified 4.7 million cis-mQTLs affecting 121.6K methylation probes, roughly 25% of Illumina Infinium HumanMethylation450K array. Bonder et al. (Bonder et al., 2017) found 272,037 independent cis-mQTLs covering 34.4% of all 405,709 CpG sites tested. More recently, a meta-analysis (Min et al., 2021) using linkage disequilibrium (LD) clumping identified 248,607 independent cis-mQTL associations accounting for 45% of the 450K array. To accurately assess the functionality of an mQTL, a more nuanced understanding of DNA methylome is needed to better model the proximity between the candidate variant and the differentially methylated CpG sites.

To address these concerns, we aim to identify a novel sequencing approach to detect DNA base modifications and genetic variations at the single-molecule level. Ultimately, we decide to leverage nanopore long-read technology to develop a more robust framework to elucidate the allele specific methylation in the human genome. Using this technology, we have directly identified 5mC and 5hmC base modifications with single-nucleotide resolution over kilobases-long reads. This technology enables PCR-free genome-scale observations of the methylome over unprecedented genomic distances. With the shift from linear regression models to directly comparing differentially methylated regions between haplotypes, the requirements for sample size have become less stringent. Moreover, the continuous 5mC profiling with nanopore technology enhances the de-novo identification of differentially methylated regions (DMRs), providing more accurate insights into the functionality of target regions and benefiting the interpretation of their underlying biological significance.

In this project, we explored the potential of nanopore long-read technology for profiling the methylome of the human genome and developed a computational pipeline to detect allele-specific methylation using adaptive sampling in prostate cells. We demonstrated the pipeline’s effectiveness in identifying functional variants and methylated regions.

## Methods

### Nanopore Open Data retrieval

Adaptive sampling (AS) is an advanced technique used in nanopore sequencing technology to selectively enrich or deplete specific regions of interest from a genomic sample in real-time. To evaluate the adaptive sampling performance, we accessed the Reduced Representation Methylation Sequencing (RRMS) result from Oxford Nanopore Open Data S3 bucket (s3://ont-open-data/rrms_2022.07/), which aimed to target 310 Mb of the human genome regions highly enriched for CpG sites, including ∼28,000 CpG islands, ∼50,600 shores and ∼42,700 shelves as well as ∼21,600 promoter regions (https://community.nanoporetech.com/attachments/7599/download). The Binary Alignment Map (BAM) files were retrieved using the AWS command line interface. Since the BAM file included alignment information to the hg38 genome and per-read base modification predictions for CpG context 5mC call, we directly used it for MONOD2 co-methylation analysis.

### Co-methylation analysis of nanopore long-reads with MONOD2

MONOD2 is a toolkit for co-methylation analysis in bisulfite sequencing data. To fit the nanopore base modification information to MONOD2 requirement, we developed a PERL script (*MMMLparse*.*pl*) to generate a bisulfite emulative BAM file based on the base modification tag in aware of the indels, mismatch and soft-clip bases in the read. After the in-silico bisulfite conversion, we generated methylation haplotype retrieving PERL script (*getHaplo_SE_cgOnly*.*pl*) for single-end read adapted from MONOD2 script (*getHaplo_PE_cgOnly*.*pl*). After the methylation haplotype pattern retrieval, the methylation haplotype block (MHB) was defined with *cghap2mhbs*.*sh* MONOD2 script. Since the co-methylation profiling estimation was based on linkage disequilibrium, we also applied a minimal methylation frequency of 30% and a minimal read depth 50 for calculating the R^2^ to avoid the bias caused by low variability site or poor read coverage. For MHB determination, we used minimal R^2^ of 0.05 to visualize all possible co-methylation sites in Figure 1B-1C and used the same R^2^ of 0.5 for direct comparison with Guo’s result (Guo et al., 2017) in Supplementary Figure 1B-1E. After the MHB determination, we used “*smoothScatter*” function from the R graphics package to visualize the distribution of R^2^ and the distance for those CpG sites in the same MHB.

**Figure 1.**
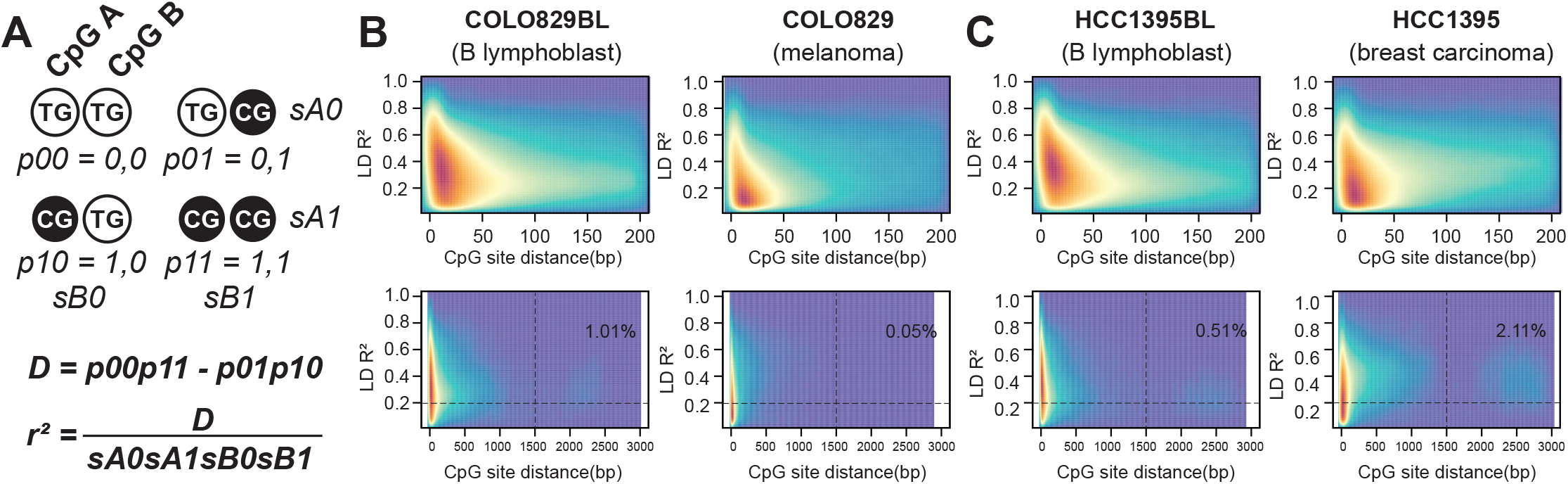
**A**. Explanation of using linkage disequilibrium R^2^ calculation for co-methylation analysis. Smooth scatter plot of co-methylation profiles for CpG sites located in the methylation haplotype block (MHB) in the normal-tumor pair of COLO829BL and COLO829 (**B**) cells, and HCC1395BL and HCC1395 (**C**) cells.

### 5mCG methylation profiling near transcription factor binding motifs

To characterize the 5mCG methylation levels surrounding known transcription factor binding sites, we used the modkit pileup function (https://github.com/nanoporetech/modkit) to generate a bedGraph file for each CpG site and then convert to bigwig file with bedGraphToBigWig program (https://www.encodeproject.org/software/bedgraphtobigwig/). We downloaded the Homo Sapiens (hg38) Familial binding site collection from the JASPAR database and generated a control BED file for each motif from the human genome with BEDtools (Quinlan and Hall, 2010). We then used deeptools (Ramirez et al., 2016) to compute and visualize cumulative methylation to each motif and its surrounding genome. From the intermediate signal matrix, we extracted the cumulative methylation percentage from the most adjacent bin to compare the transcription factor (TF) motif region and the control region in t-tests.

### Target design of adaptive sampling (AS)

After evaluating the performance of nanopore RRMS demo data, we aimed to design a customized AS BED file to enrich sequences at GWAS and mQTL variant loci. We downloaded prostate cancer risk variants reported by the GWAS catalog (https://www.ebi.ac.uk/gwas/home). We then used LDlink (Machiela and Chanock, 2015) to retrieve variants highly associated with the tag SNPs (LD R^2^ ≥ 0.5) in either of the populations below: African (AFR), Admixed American (AMR), East Asian (EAS), European (EUR) or South Asian (SAS). As a result, we obtained a total of 68,436 SNPs with prostate cancer risk association.

We also downloaded mQTL summary statistics of TCGA prostate cancer cohort (Gong et al., 2019) and filtered the association by the regression coefficient (R ≥ 0.5 or ≤ -0.5). As a result, we found 137,825 SNPs with strong mQTL effect. To generate an AS BED file, we expanded 4000 bp on both sides of each SNP locus and merged the expanded intervals. As an outcome, the BED file covered 250,689,518 bp with 5,569 intervals, roughly 8.36 % of the human genome.

### AS nanopore sequencing in prostate cell 22Rv1

The 22Rv1 (RRID: CVCL_1045) cell was obtained from the ATCC and grown in RPMI1640 medium supplemented with 10% fetal bovine serum (FBS). After harvesting the cell during the log phase, we used Puregene Cell Kit (QIANGEN, 158043) to extract high-quality genomic DNA with RNase A treatment included. The genomic DNA was sheared into 8 kb fragments with g-TUBE (Covaris, 520079). The sheared DNA was used for PCR-free nanopore sequencing library preparation using Ligation Sequencing Kit (Oxford Nanopore Technology, SQK-LSK110). After library preparation, 100 fmol library solution was loaded to R9.4.1 flow cell. With the target BED file and the matched reference FASTA plugged in, we started the data acquisition in MinKNOW and kept running until the flow cell efficiency was fully depleted.

### Nanopore adaptive sampling sequencing base-calling and read-level phasing

After the sequencing finished, we used dorado basecaller on NVIDIA GPU to infer DNA sequence and CpG site base modification based on the model “dna_r9.4.1_e8_sup@v3.3_5mCG_5hmCG”. The base-calling step generated hg38 alignments with per-read base modification information for downstream phasing and methylation analysis. To evaluate the enrichment performance of the AS run, we used the “mosdepth” program (Pedersen and Quinlan, 2018) to summarize the coverage for the genome and the target region. To resolve the SNP calling difficulty in the homopolymer region and improve the accuracy of SNP calling for nanopore long read data, we applied a pipeline called “PEPPER-Margin-DeepVariant” (Shafin et al., 2021) to provide the state-of-the-art variant calling for the 22Rv1 genome. We then used the candidate variants for read-level phasing and haplotype tagging in longphase (Lin et al., 2022).

### Haplotype-specific methylation calling and de-novo differential methylated region (DMR) analysis

To generate multiple observations for the same CpG site, we used modkit pileup function to call 5mCG methylation with haplotype-partitioning and strand specific output. The output bedGraph files were used to identify de-nono DMRs between the two haplotypes in “metilene” software. To improve sensitivity and fit better with the long read sequencing data, we increased the “--maxdist” parameter (allowed distance between two CpGs within a DMR) to 1000 bp and lower the “--mincpgs” parameter (minimum number of CpGs in a DMR) to 5.

### Correlation between chromatin accessibility and CpG methylation

To quantify the chromatin accessibility in the 22Rv1 cell, we calculated the Tn5 insertion coverage by normalizing the Transposase cut event count in each peak to the peak width. We also calculated the averaged methylation percentage for each peak. The paired profiling was plotted with the “smoothScatter” function from the R graphics package for visualization.

### Data accession and source code repository

The source code generated in this study can be accessed through the GitHub repository (https://github.com/Yijun-Tian/Nanopore). The 22Rv1 ATAC-seq data was obtained from the GEO accession GSE264518 (Tian et al., 2024). The eQTL data of prostate tissue was retrieved from GTEx Consortium (Consortium, 2020). The prostate cancer mQTL data was downloaded from Pancan-meQTL database (Gong *et al*., 2019). The raw data for 22Rv1 sequencing was stored with the GEO accession number PRJNA1158791 (https://dataview.ncbi.nlm.nih.gov/object/PRJNA1158791?reviewer=ovnrt2bnad9muc43sjsa2nc5os).

## Results

### Co-methylation profiling guided by long-read sequencing

The co-methylation analysis intends to study the correlation between different CpG sites to identify genomic regions showing coordinated epigenetic changes. With bisulfite-emulated nanopore data, we used the LD R^2^ from the MONOD2 method to measure the non-random association of the methylation status between two CpG sites (**Figure 1A**), in which an R^2^ value close to 1 indicates strong co-methylation, while a value near 0 indicates random methylation. In the RRMS data (**Figure 1B, upper panel**), we visualized the LD R2 profiling with the CpG site distance and found that most of the LD R^2^ in cancerous cell lines (COLO829 and HCC1395) was between 0 to 0.2, while the value in parental lymphoblast cell lines (COLO829BL and HCC1395BL) was between 0.2 to 0.6, suggesting an overall diminished synchronization of the epigenetic profiling in cancer cells. The long read (**Figure 1B, lower panel**) profiling also demonstrated a small proportion of distant CpG sites (≥1500 bp) showed weak-to-median co-methylation (R^2^≥0.2), suggesting an operation of unison between these sites. As an example (**Supplementary Figure 1A**), we demonstrated a locus with the long-range co-methylation on chromosome 13 in the 4 samples. According to Guo et al., we also applied a more stringent criterion of R^2^ for MHB identification to see whether those highly correlated sites exsist in the nanopore data. However, with a minimal R^2^ of 0.5, we did not find the perfect coupled sites (**Supplementary Figure 1B-1E**) as observed in the bisulfite sequencing data (Guo *et al*., 2017).

### CpG methylation footprint of human transcription factor (TF)

With the comprehensive AS coverage on the CpG dense region, we were able to estimate the epigenetic profiling in the genome surrounding the known human TF motifs. We scanned a total of 835 TF binding motifs and identified 251, 269, 194, and 148 TFs with significantly altered methylation (p ≤ 0.05) in COLO829BL, COLO829, HCC1395BL, and HCC1395 cell line, respectively. In COLO829BL and COLO829 cell lines, we highlighted CTCF (**Figure 2A**), NRF1 (**Figure 2B**) and ZNF93 (**Figure 2C**) as the TF associated with hypomethylation in the nearby genome, and RARA (**Figure 2D**) as the TF associated with hypermethylation. Consistently, we observed similar trends in HCC1395BL and HCC1395 cells (**Supplementary Figure 2A-2D**).

**Figure 2.**
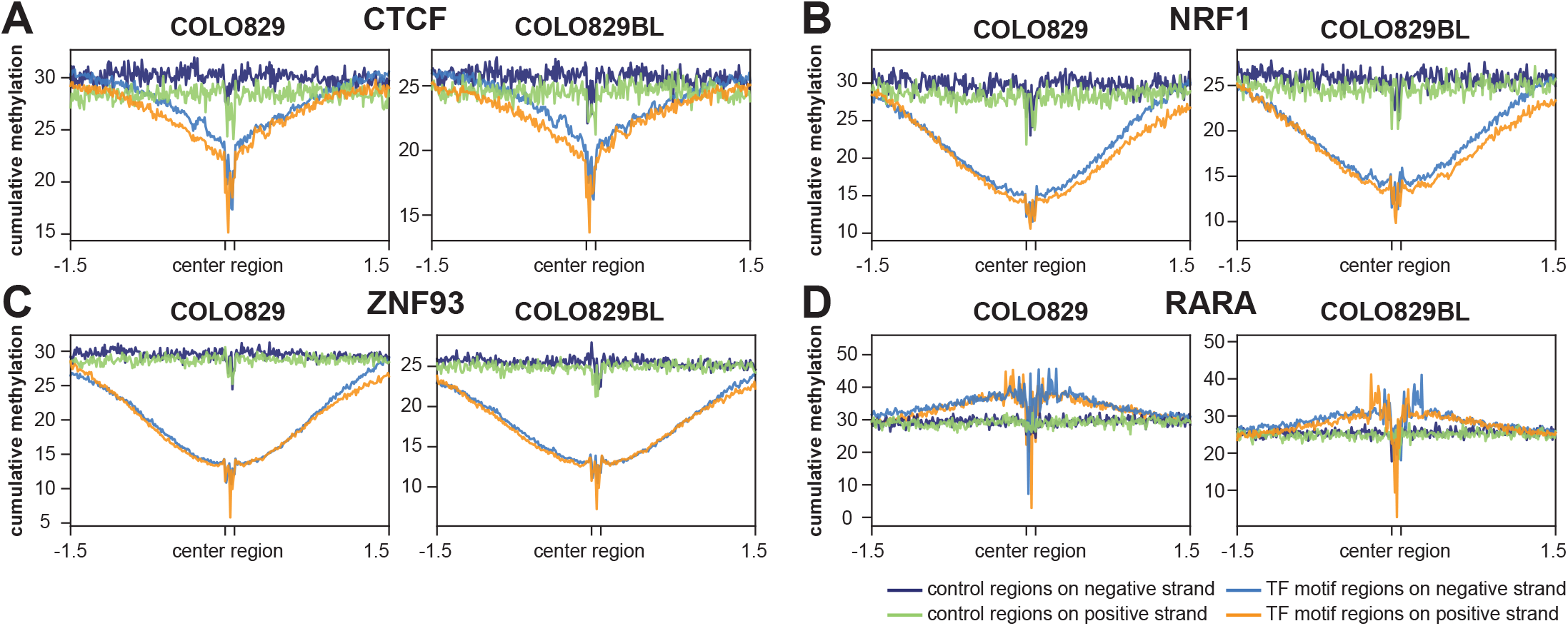
In COLO829BL and COLO829 cells, CpG methylation status surrounding known transcription factor binding motifs, such as CTCF (**A**), NRF1 (**B**), ZNF93 (**C**) and RARA (**D**).

### Adaptive sampling nanopore sequencing targeting phenotypical variant

To selectively observe the DNA methylation patterns related to germline risk variants linking to prostate cancer phenotypes, we examined GWAS catalogue (https://www.ebi.ac.uk/gwas/home) and TCGA prostate cancer mQTL databases and identified a total of 68,436 SNPs with prostate cancer risk association and 137,825 SNPs with strong mQTL effect (regression coefficient R ≥ 0.5 or ≤ -0.5) (**Figure 3A**). Interestingly, we only found 2,539 SNPs overlapped between the two groups. Compared to genome background, the AS enriched 3.88 and 4.76 folds more sequences at mQTL and GWAS SNP regions, respectively (**Figure 3B**). Across the 22 chromosomes, we found a significant negative correlation between the target length (proportion of target region length to the belonging chromosome) and the fold of enrichment (**Figure 3C**).i Through grouping the methylation call by their location, we further identified expected hypomethylation for CpG sites located in CpG island (**Figure 3D**), and hypermethylation for CpG sites located outside CpG island (**Figure 3E**). To find haplotype-specific methylated regions, we performed de-novo DMR analysis between the two haplotypes of the 22Rv1 genome, treating the methylation levels reported by each DNA strand as replicates since most 5mCG exist symmetrically at CpG dinucleotides. The DMR analysis found 6,390 genomic locations showed haplotype-specific 5mCG modification in the 22Rv1 genome (p-MWU ≤ 1e-5 and delta ≥ 50%). With the help of the “genomicDensity” function, we identified apparent imprinting regions consisting of DMR clusters across the genome in the circle plot (**Figure 3F**). The full list of DMRs can be accessed in **Supplementary Table 1**. Furthermore, we also found that some DMRs are likely caused by allele-specific methylation, offering strong function hints for the target variant.

**Figure 3.**
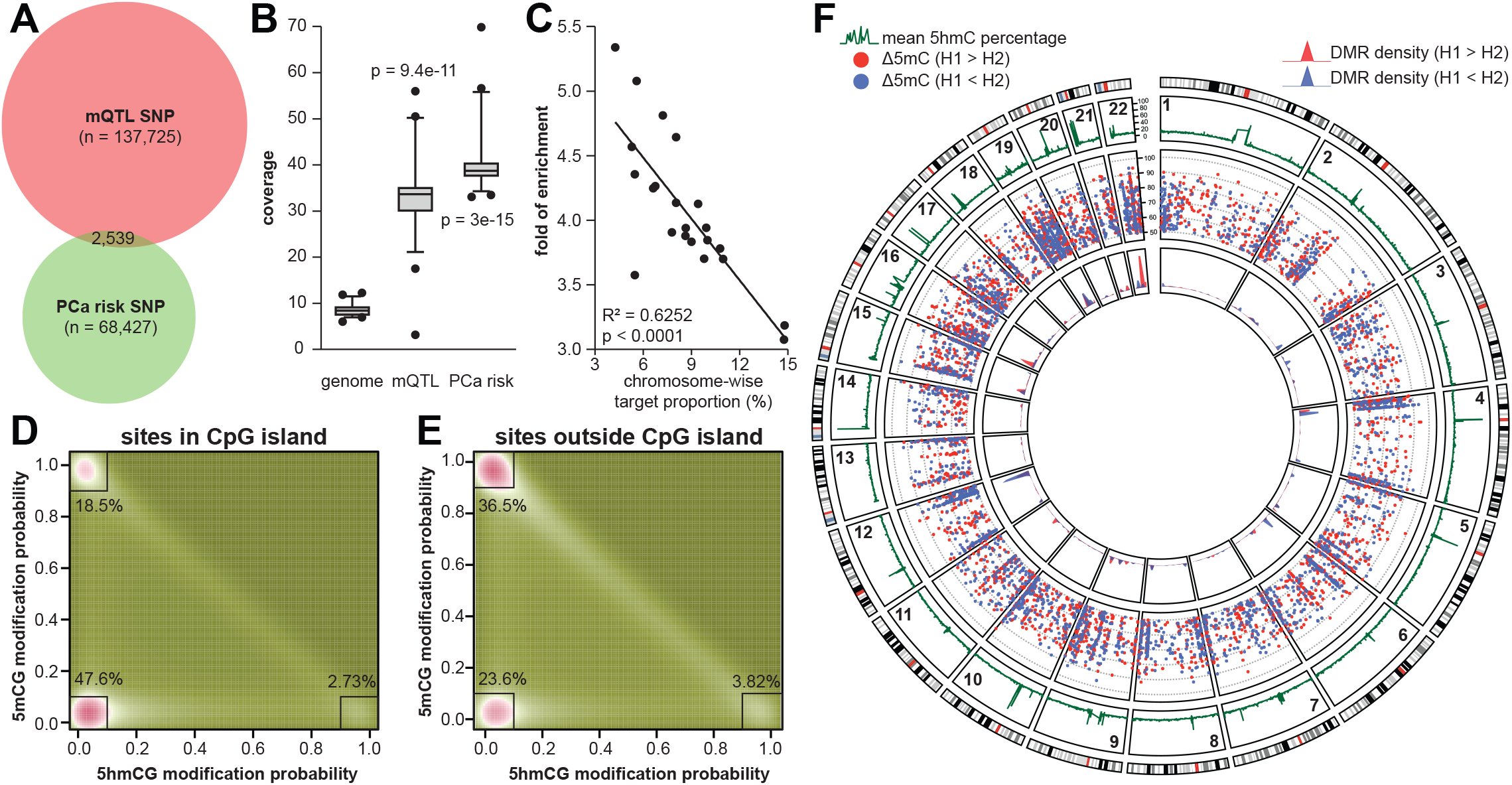
**A**. Proportional Venn diagram of the target SNP with TCGA mQTL signals and GWAS risk significance. **B**. Sequencing coverage summarized by chromosomes with adaptive sampling run in 22Rv1 genome. **C**. Correlation between the target length and enrichment fold across the 22 autosomes. Distribution of dual methylation calling for the CpG sites in (**D**) or outside (**E**) CpG island in chromosome 19. The percentages highlight the proportion of highly confident modified sites within the nearby squares. **F**. Circular plot of 5hmC modification percentage (green line, outermost circle), DMR methylation difference (red and blue scatters, middle circle), and the DMR density (red and blue peaks, innermost circle).

### Allele-specific methylation associated with GWAS or mQTL variant

Previously, we reported a functional prostate cancer risk variant causing allele-specific methylation in the 22Rv1 genome with bisulfite amplicon sequencing (Tian *et al*., 2022). In nanopore sequencing data, we found that the risk SNP rs7247241 was adjacent to a 162-bp co-methylated region (chr19:38256705-38256867), with the T allele consistently hypermethylated than the C allele (**Figure 4A**). Additionally, the ATAC-seq result demonstrated that the hypomethylated C allele tended to be more accessible than the hypermethylated T allele (**Figure 4A**). Although no mQTLs are being reported for this locus, we found that the risk allele (T) was significantly associated with elevated PPP1R14A gene expression (**Figure 4B**), identified as an oncogene in our previous work. We also highlighted another locus mapping to the KCNIP3 gene, with the genotype of mQTL SNP rs2113417 associated with the methylation level of a 710-bp region (chr2:95333760-95334470) (**Figure 4C**). Consistently, we identified an even stronger allele-specific chromatin accessibility in the region containing rs2113417 (**Figure 4C**). Interestingly, of the three mQTLs reported for rs2113417, only CpG site cg03595348 is located within the same DMR as the SNP, although its p-value did not show the best (**Figure 4D**). To describe the relationship between the chromatin accessibility and the methylation level at the genome scale, we calculated Tn5 insertion coverage and mean methylation percentage for each ATAC-seq peak, and found that two profiles separated the peaks into two main clusters, with the major one showing high methylation (∼80%) and low chromatin accessibility, and the minor one showing low methylation (less than 10%) and high chromatin accessibility (**Figure 4F**).

**Figure 4.**
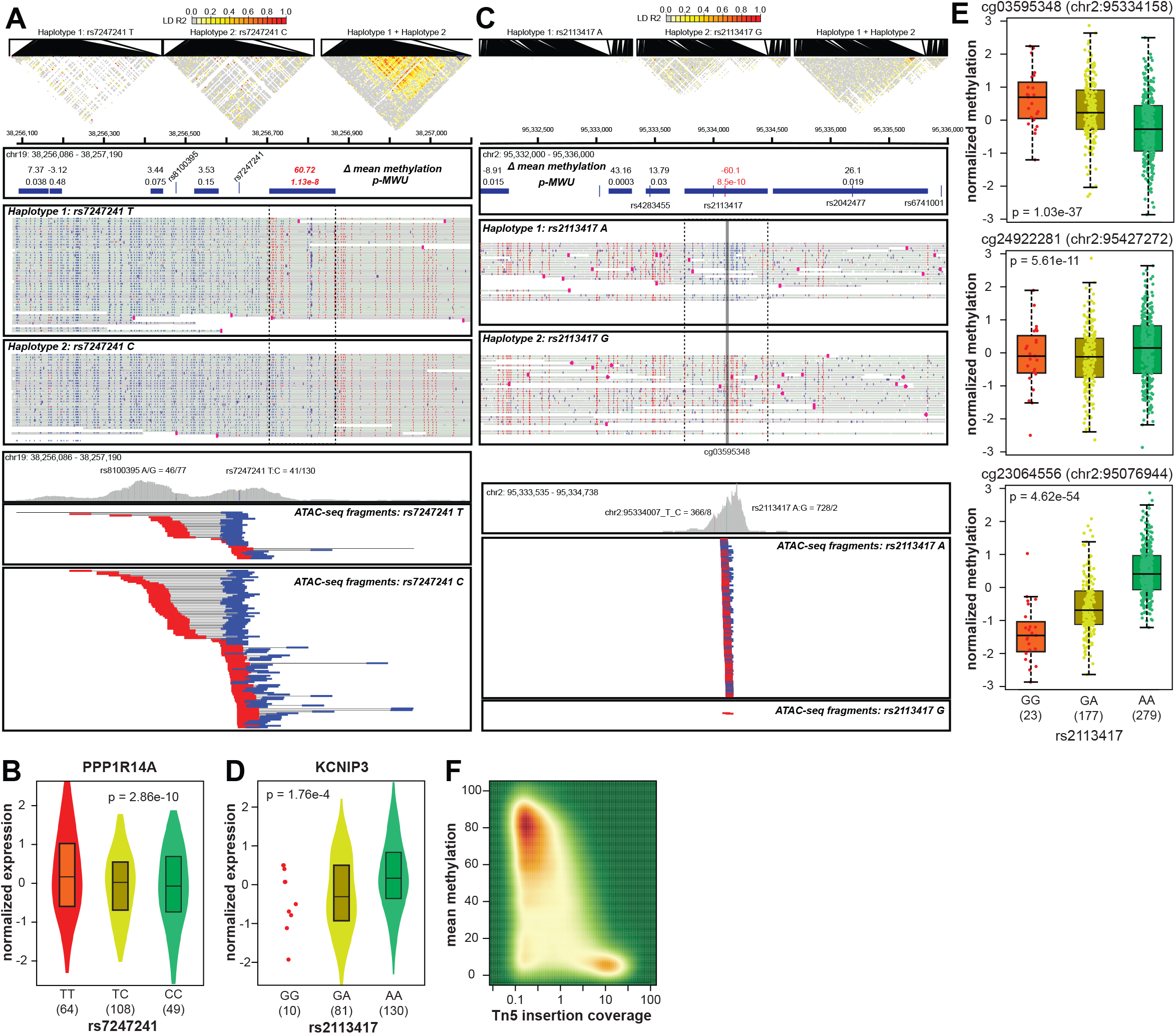
**A**. Exemplary differential methylated region (DMR) identified near the GWAS risk variant rs7247241 and the chromatin accessibility profiling with the rs7247241 alleles in 22Rv1 cells. **B**. GTEx eQTL results for rs7247241 genotype and PPP1R14A expression in prostate tissue. **C**. Exemplary differential methylated region (DMR) identified near the mQTL variant rs2113417 and the chromatin accessibility profiling with the rs2113417 alleles in 22Rv1 cells. **D**. GTEx eQTL results for rs2113417 genotype and KCNIP3 expression in prostate tissue. **F**. TCGA mQTL results for rs2113417 in prostate cancer tissue.

## Discussion

The technical concern of bisulfite sequencing has been thoroughly discussed over the last decade, particularly regarding its potential to fragment DNA templates and introduce PCR amplification bias during sample preparation. In the co-methylation analysis based on WGBS with short-read NGS, Guo et al. (Guo *et al*., 2017) identified that over 80% of those CpG sites located in the same methylation haplotype block were with high LD (R^2^ ≥ 0.9), regardless of the cell type. However, in a PCR-free condition of nanopore long-read sequencing, we could not identify these highly associated co-methylation sites. One of the hypotheses is that during the PCR amplification step, co-methylated molecules may have been over-amplified, potentially introducing artificial LD sites. Another possibility is that when applying the population-based statistic to evaluate read-level co-methylation status, several additional metrics must be considered, such as the minimum read depth and minor allele frequency of the two CpG sites. According to previous discussion (Eberle et al., 2006; VanLiere and Rosenberg, 2008; Wray, 2005), R^2^ calculation is dependent on allele frequencies of the two loci, which could easily generate perfect LD (R^2^ = 1.0) when both CpG sites are with extremely high and low methylation frequencies. Therefore, we applied a minimal read depth of 50 and a minor allele frequency of 0.3 for the co-methylation R^2^ calculation, aiming to focus on those sites of high confidentiality in the PCR-free condition.

The long-read technology also provided a broader vision to study epigenetic co-regulation. Ideally, sequencing the human genome at the telomere-to-telomere scale would reveal a full spectrum of possible long-range interactions based on epigenetic modification. However, the technical issue is that as the read length increases, the nanopore sequencing throughput are reduced due to the complex DNA secondary structure or heavy modification genomic region obstruction, which explains why the RRMS demo data used 8kb fragmented DNA for maximum read coverage. Based on the observations of the distant co-methylation (**Supplementary Figure 1A**), the application successfully reaching high coverage with extended read lengths may provide greater insight into these longer-range interactions. Our analyses of the methylation profiling on known binding motifs also described a unique epigenetic footprint for human TFs. The observation of methylation changes of CTCF motif is consistent with previous studies (Damaschke et al., 2020), suggesting that the CTCF is a major in directing localized DNA hypermethylation. Other TFs reported to be sensitive to or associated with DNA methylation, such as NRF1 (Domcke et al., 2015), ZNF93 (Fukuda et al., 2022) and RARA (Hassan et al., 2017), also demonstrated consistent adjacent methylation changes in our analysis.

With the increasing availability of powerful GPU resources, more and more applications use nanopore adaptive sampling (AS) to enrich target-of-interest or deplete unwanted genome (Kovaka et al., 2021; Loose et al., 2016). The computational approach utilizes GPU-based fast basecalling to perform raw electrical signal mapping to make real-time “Read Until” decisions to select specific DNA molecules. Here, we successfully applied the AS approach to enrich the germline variant regions, aiming to obtain high PCR-free coverage to identify allele-associated epigenetic changes. However, due to the relatively high error rate in nanopore reads, the alignments are frequently filled with small indels, especially in the homopolymer regions, which poses significant challenges for the subsequent variant calling and haplotype phasing analysis. To this end, we applied a neural network tool called DeepVariant (Poplin et al., 2018; Shafin *et al*., 2021) to polish the candidate variants to achieve the best calling accuracy. Based on the continuity nature of CpG methylome, we implemented a de-nono DMR selection procedure to identify clusters of CpG sites that showed modification differences between the two haplotypes. Considering that the haplotype-specific DMRs don’t inform the potential causal variants, we included additional information of the closest phasing SNP to each DMR and also annotated further whether the SNP is an GWAS or mQTL variant. From the results, we identified two candidate variants with known or potential functions in prostate cells. For example, the risk variant rs7247241 has been found to be associated with nearby genome methylation in bisulfite sequencing, and our DMR analysis reproduced this finding. For another SNP rs2113417, we successfully applied the nanopore data to discover the most related DMR between the alleles, which further revealed that the biological origin is from the CpG site cg03595348 among the mQTL signals. This result suggests that the haplotype-specific DMR approach may help remove the mQTL redundancy and improve causal association discovery. Additionally, considering the minor allele frequency of rs2113417, the nanopore DMR approach also provides valuable information for observing the epigenetic effect driven by those rare alleles. One shortcoming of using the utility to a population cohort is that the read-level variant calling and phasing based on the nanopore long-read data is currently limited to each diploid individual, which will generate different haplotype phase set information according to different persons. This could challenge the between-haplotype DMR identification. To further expand this utility to a multiple-sample design, a genotype-based reference would be helpful for further harmonizing and standardizing the haplotype information so that all samples could be compared and analyzed together.

Taken together, we spotlighted multiple advantages of the nanopore long-read sequencing for methylation analysis and envisioned an innovative utility with this technology to address the technical issue in determining allele-specific methylation. The analytical framework introduced in this study for long-read DNA methylation analysis offers a new perspective and uncovers previously unknown insights in the field of genetics.

## Acknowledgements

This study has been supported by the National Institutes of Health [R01CA250018 and R01CA212097, to L.Wang. and R01CA263494 to L.Wu]. The funders had no role in study design, data collection and analysis, publication decision, or manuscript preparation.

## Author Contributions Statement

Conception and design: Y. Tian, S. McDonnell, N. Larson, L. Wu, L. Wang Development of methodology: Y. Tian, S. McDonnell, N. Larson, L. Wu, L. Wang Acquisition of data: Y. Tian

Analysis and interpretation of data: Y. Tian, L. Wu, S. McDonnell, N. Larson

Writing, review, and revision of the manuscript: Y. Tian, S. McDonnell, N. Larson, L. Wu, L. Wang Study supervision: Y. Tian, N. Larson, L. Wu, L. Wang

---

**supplementary figure 1.**
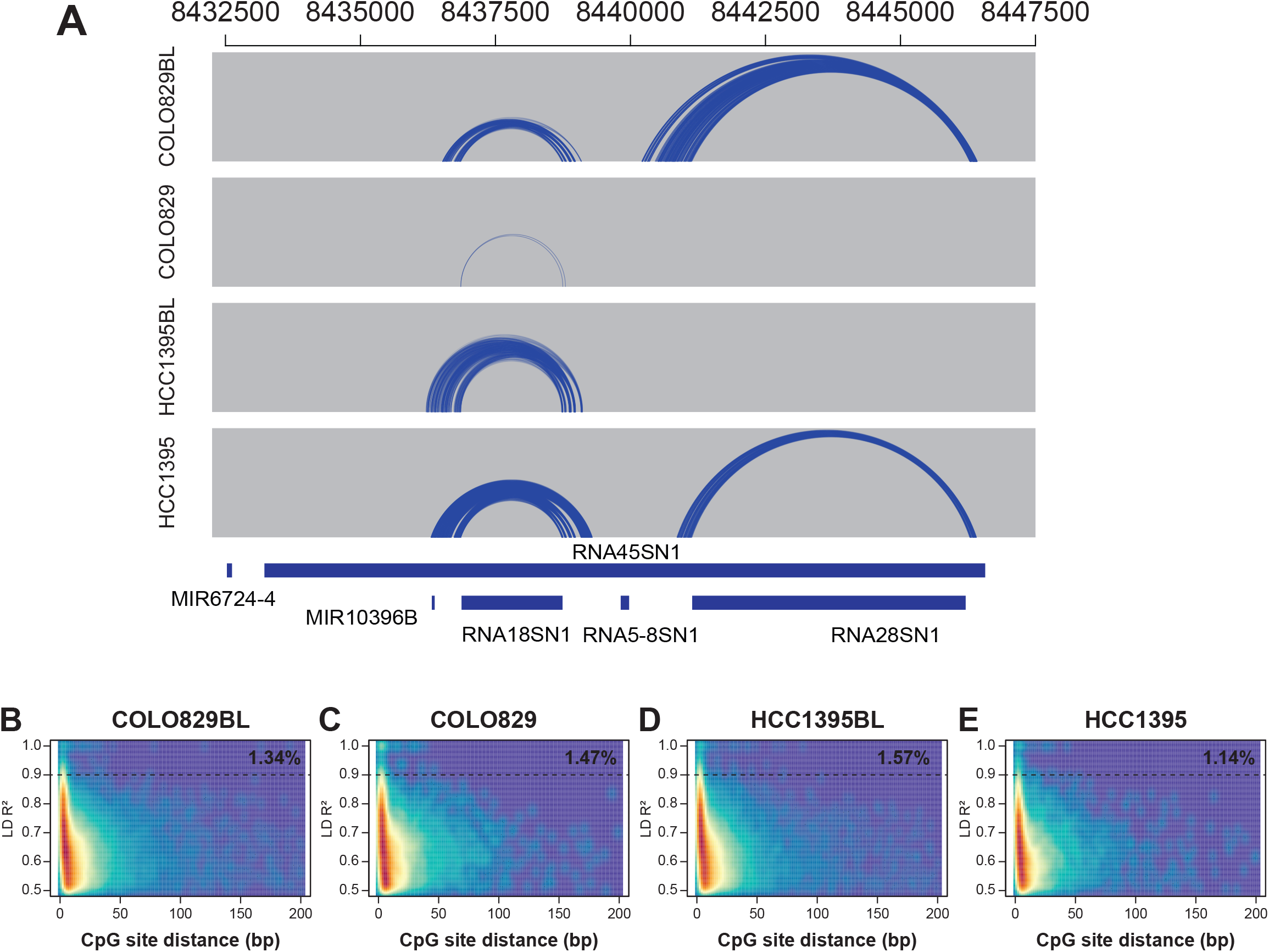

**supplementary figure 2.**
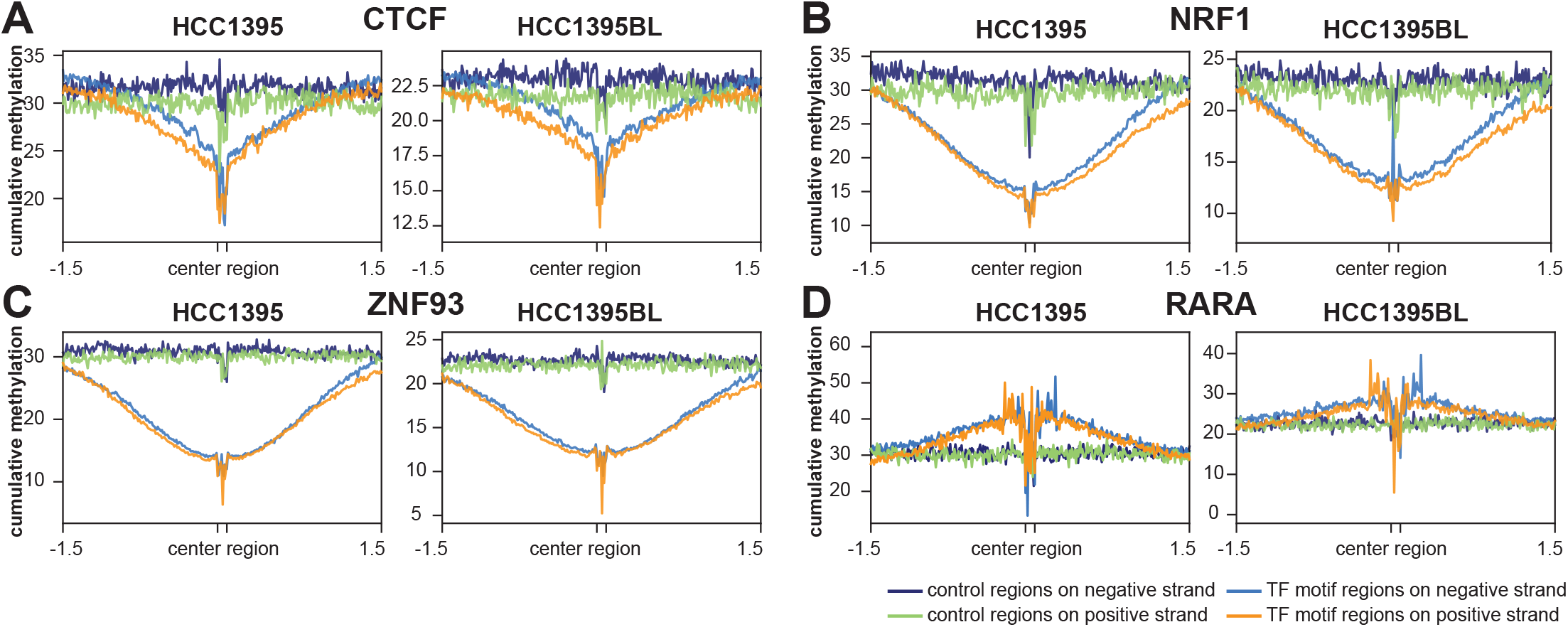

